# Host nutrition mediates interactions between plant viruses, altering transmission and predicted disease spread

**DOI:** 10.1101/761254

**Authors:** Amy E. Kendig, Elizabeth T. Borer, Emily N. Boak, Tashina C. Picard, Eric W. Seabloom

**Affiliations:** Department of Ecology, Evolution, and Behavior, University of Minnesota, St. Paul, MN 55108, USA; Agronomy Department, University of Florida, Gainesville, FL 32611, USA; Department of Horticultural Sciences, Texas A&M University, College Station, TX 77843, USA

**Author notes:** Corresponding author: Amy Kendig.

**Keywords:** co-infection, transmission, within-host, disease spread, barley and cereal yellow dwarf viruses, *Avena sativa*, nitrogen, phosphorus, *Rhopalosiphum padi*

## Abstract

Interactions among co-infecting pathogens are common across host taxa and can affect infectious disease dynamics. Host nutrition can mediate these among-pathogen interactions, altering the establishment and growth of pathogens within hosts. It is unclear, however, how nutrition-mediated among-pathogen interactions affect transmission and the spread of disease through populations. We manipulated the nitrogen (N) and phosphorus (P) supplies to oat plants in growth chambers and evaluated interactions between two aphid-vectored Barley and Cereal Yellow Dwarf Viruses: PAV and RPV. We quantified the effect of each virus on the other’s establishment, within-plant density, and transmission. Co-inoculation significantly increased PAV density when N and P supplies were low and tended to increase RPV density when N supply was high. Co-infection increased PAV transmission when N and P supplies were low and tended to increase RPV transmission when N supply was high. Despite the parallels between the effects of among-pathogen interactions on density and transmission, changes in virus density only partially explained changes in transmission, suggesting that virus density–independent processes contribute to transmission. A mathematical model describing the spread of two viruses through a plant population, parameterized with empirically derived transmission values, demonstrated that nutrition-mediated among-pathogen interactions could affect disease spread. Interactions that altered transmission through virus density–independent processes determined overall disease dynamics. Our work suggests that host nutrition alters disease spread through among-pathogen interactions that modify transmission.

## Introduction

Resource supply can alter the outcome of species interactions (Tilman 1977, Maestre and Cortina 2004). A rich body of theoretical and empirical literature has explored the effects of resource supply on ecological dynamics, but most has focused on free-living organisms (Bruno et al. 2003, Miller et al. 2005). The nutrients consumed by hosts (i.e., host nutrition) are important mediators of resource supply to assemblages of pathogens and other symbionts (Smith et al. 2005). Host nutrition can affect interactions among pathogens that co-infect plant or animal hosts (Lacroix et al. 2014, Lange et al. 2014, Budischak et al. 2015, Wale et al. 2017)— interactions that can influence host survival, transmission between hosts, and evolution of virulence (Vasco et al. 2007, Tollenaere et al. 2016). Therefore, host nutrition may affect infectious disease dynamics by altering among-pathogen interactions.

Among-pathogen interactions can have positive, neutral, or negative effects on within-host pathogen fitness (Moreno and López-Moya 2020). Competition among pathogens for limiting resources, such as nutrients, cells, or tissues, can suppress pathogen densities (Smith and Holt 1996, Pedersen and Fenton 2007). Host nutrition that affects the supply of pathogen-limiting resources can alter the outcome of pathogen competition (Wale et al. 2017). Pathogens also can interact indirectly by promoting or suppressing host immune reactions (Pedersen and Fenton 2007, Vasco et al. 2007). Immune functioning in mammals depends on vitamins, zinc, iron, and iodine (Katona and Katona-Apte 2008), and plant susceptibility to infection can depend on nitrogen (N), phosphorus (P), and potassium (K) in the soil (Dordas 2009). It follows that host nutrition also can affect immune-mediated pathogen interactions (Budischak et al. 2015).

Interactions among pathogens can affect disease spread when there is a strong relationship between within-host pathogen density and processes that affect host population dynamics, including transmission, mortality, and recovery (Mideo et al. 2008, Handel and Rohani 2015). Pathogens that reach higher densities within hosts are more likely to produce more propagules for transmission (McCallum et al. 2017). Interactions among pathogens within plants and animals alter transmission and the proportion of the population that becomes infected (Ezenwa and Jolles 2011, Susi et al. 2015a, Halliday et al. 2017). Yet, it is unclear how the impacts of host nutrition on among-pathogen interactions affect disease spread. Nutrition-mediated interactions within the host are likely to influence disease spread if a strong relationship between within-host pathogen density and a process that affects host population dynamics (e.g., transmission) exists (Gilchrist and Coombs 2006, Strauss et al. 2019).

Increases in within-host pathogen densities do not always increase the probability of transmission (Handel and Rohani 2015, McCallum et al. 2017). For example, the relationship between pathogen density and transmission is sigmoidal for malaria-inducing *Plasmodium falciparum*, and increases in *P. falciparum* density beyond a threshold do not affect transmission (Alizon and van Baalen 2008). Such non-linearities can arise when vector behavior and pathogen-vector interactions affect the probability of transmission (Gray et al. 1991), decoupling transmission from within-host dynamics. Interactions among pathogens also can change establishment or transmission independently of changes in pathogen density. For example, co-infection can modify vector preference and the efficacy of vector transmission, causing transmission from co-infected hosts to differ from singly infected hosts (Rochow et al. 1983, Srinivasan and Alvarez 2007). Therefore, nutrition-mediated among-pathogen interactions may modify disease spread through processes that are independent of within-host density.

Here we experimentally tested the effects of host nutrition on among-pathogen interactions and transmission using a well-studied group of aphid-transmitted viruses that infect crops and wild plants: the Barley and Cereal Yellow Dwarf Viruses (B/CYDVs; Power et al. 2011). In a growth chamber experiment, we manipulated soil N and P concentrations supplied to oat plants singly- and co-inoculated with two B/CYDVs: BYDV-PAV (PAV, hereafter) and CYDV-RPV (RPV, hereafter). We quantified the effects of interactions between the viruses by measuring establishment, within-plant virus densities, and transmission to new plants (Fig. 1). Then, we used empirically estimated transmission values to parameterize a mathematical model with the goal of predicting the effects of host nutrition-mediated among-pathogen interactions on disease spread. We used this experiment and model to address the following questions.

**Figure 1.**
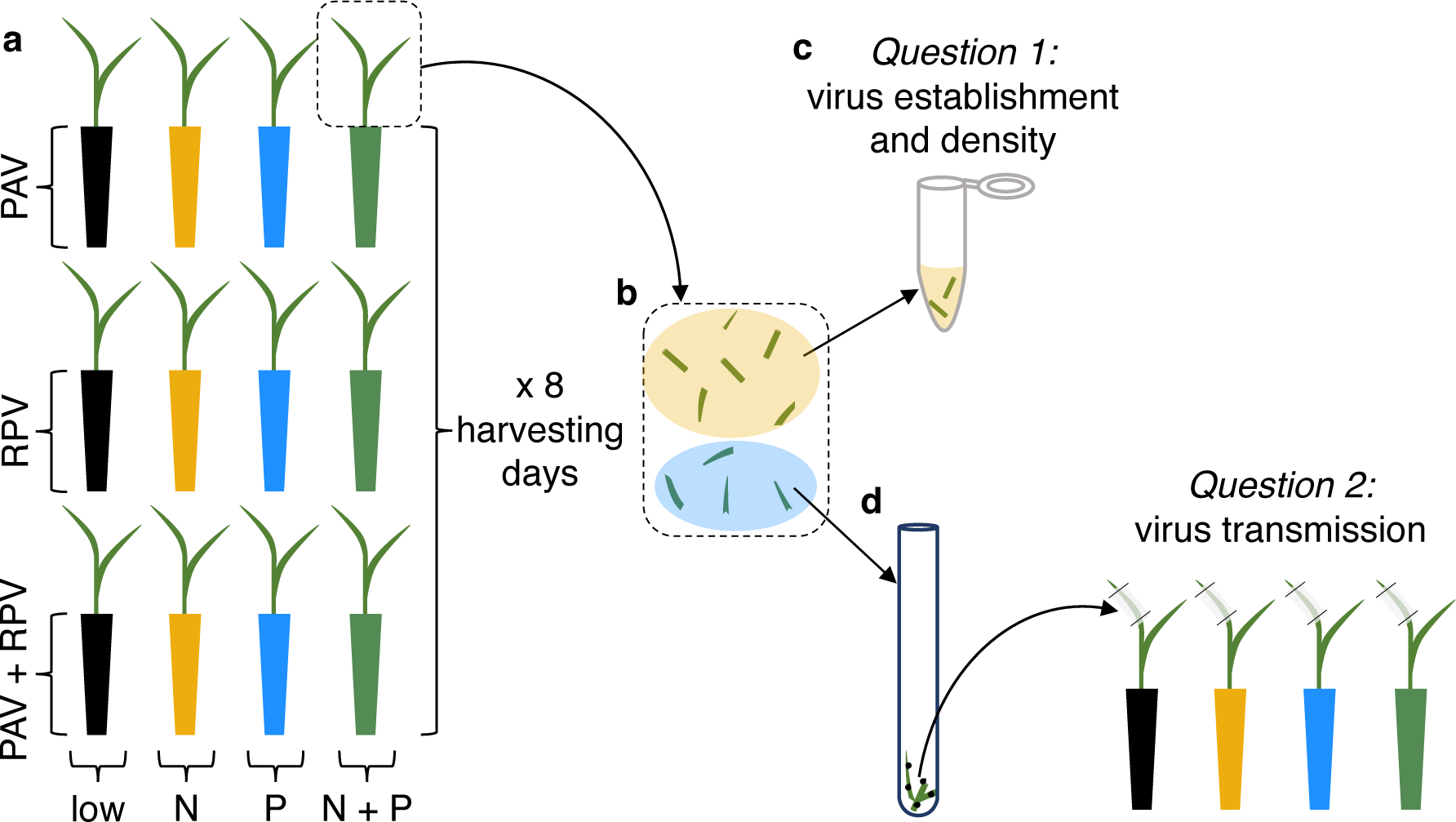
Diagram of the experimental design. (a) Source plants were grown in one of four nutrient treatments and inoculated with one of three inoculation treatments. (b) Source plants were harvested at eight different time points and tissue was cut into pieces. (c) Tissue was used in molecular analysis to determine virus establishment and density. (d) Tissue was placed in tubes with aphids, which were used to inoculate recipient plants and assess transmission.

### Question 1: Does host nutrition affect among-pathogen interactions within hosts?

Results from previous studies indicate that host nutrition mediates B/CYDV replication and among-pathogen interactions (Lacroix et al. 2014, Whitaker et al. 2015). However, it is unclear whether previously observed reductions in RPV infection prevalence due to co-inoculation with PAV (Lacroix et al. 2014) represent interactions that allow establishment, but suppress RPV density below detection thresholds over time, or that interfere with establishment. These two outcomes of PAV–RPV interactions could have different effects on host health and RPV transmission. Time series of virus densities can help clarify when among-pathogen interactions occur. We used a full factorial combination of high and low N and P supply rates to evaluate how host nutrition affected the nature (i.e., positive, neutral, negative) and timing (i.e., early or late relative to inoculation) of interactions between B/CYDVs within plants. While this experimental design allowed us to test whether co-inoculation altered establishment or post-establishment processes, it did not allow us to fully discern the mechanisms behind observed among-pathogen interactions (e.g., resource competition, immune-mediated interactions).

### Question 2: Does host nutrition modify among-pathogen interactions to affect transmission?

Because insect-vectored viruses with higher within-plant densities often have higher transmission (Froissart et al. 2010), among-pathogen interactions that promote (suppress) virus density are expected to promote (suppress) transmission. Virus density–transmission relationships, however, may be virus-specific (Gray et al. 1991). In addition, the impacts of co-infection on vector acquisition may affect transmission independently of virus density (Rochow et al. 1983, Wen and Lister 1991). We evaluated the effects of virus density, host nutrition, and co-infection on transmission from source plants (i.e., those inoculated in *Question 1*) to recipient plants grown with a full factorial combination of low and high N and P supply rates.

### Question 3: Can nutrition-mediated among-pathogen interactions affect disease spread?

Higher transmission can increase the rate of disease spread through a population. Thus, changes in transmission due to host nutrition-mediated among-pathogen interactions are expected to have population-level consequences. Nutrient additions have altered B/CYDV infection prevalence in wild grass populations (Seabloom et al. 2013, Borer et al. 2014), but it is unclear whether these changes were mediated by within-plant dynamics. We parameterized a two-pathogen compartmental model with our empirical results to quantify the effects of nutrition-mediated among-pathogen interactions on infection prevalence over time. We then used the model to evaluate the contribution of within-plant virus density to disease dynamics.

## Methods

### Study system

The B/CYDV group consists of single-stranded RNA viruses in the *Luteoviridae* family that can infect over 100 species of *Poaceae* and are persistently transmitted by several aphid species (Power et al. 2011). The aphid *Rhopalosiphum padi* is an effective vector of the two virus species we used in this study, PAV and RPV (Gray et al. 1991, Power et al. 2011). We maintained cultures of PAV and RPV in *Avena sativa* L. cv. Coast Black Oat (National plant germplasm system, USDA) by periodically feeding infected plant tissue to *R. padi* aphids, which were temporarily transferred to uninfected *A. sativa. Rhopalosiphum padi* colonies were maintained on uninfected *A. sativa* plants. We obtained the virus isolates from Dr. Stewart Gray at Cornell University (Ithaca, NY, USA), and the aphids from Dr. George Heimpel at the University of Minnesota (St. Paul, MN, USA), who each collected these organisms in their respective states. Uninfected plants, infected plants, and plants with *R. padi* were grown in Sunshine MVP potting soil (Sun Gro Horticulture, Agawam, MA, USA) and kept in separate growth chambers at 20°C with a 16:8 h light:dark cycle for one year prior to the experiment.

### Experimental design

The experiment was carried out from February to August 2014 over five temporal blocks. *Avena sativa* seeds were germinated in 164 mL conical pots with 70% Sunshine medium vermiculite (vermiculite and <1% crystalline silica; Sun Gro Horticulture) and 30% Turface MVP (calcined clay containing up to 30% crystalline silica; Turface Athletics, Buffalo Grove, IL, USA) that had been saturated with tap water. Beginning two days after planting, we watered each plant with one of the following modified Hoagland solutions (i.e., nutrient treatments, Appendix S1: Table S1, Hoagland and Arnon 1938): 7.5 μM N and 1 μM P (“Low”), 7.5 μM N and 50 μM P (“P”), 375 μM N and 1 μM P (“N”), or 375 μM N and 50 μM P (“N+P”), which differentially affect plant growth and B/CYDV infection prevalence (Seabloom et al. 2011, Lacroix et al. 2014, 2017). Plants were watered with 30 mL of nutrient solution twice per week prior to inoculation and weekly following inoculation. After inoculation, plants were moved to a growth chamber maintained at 20°C with a 16:8 h light:dark cycle under 28W bulbs.

We inoculated the plants used to assess pathogen establishment and density (*Question 1*) 10 to 11 days post planting with PAV, RPV, or both (Fig. 1a). Aphids fed on virus culture leaves for approximately two days and then were combined into plastic containers by inoculation type. We attached one mesh cage to each plant on the largest leaf and placed ten aphids in each mesh cage, allowing them to feed on the plants for approximately four days. PAV inoculations involved five aphids that fed on PAV-inoculated culture leaves and five that fed on uninfected leaves, RPV inoculations involved five aphids that fed on RPV-inoculated culture leaves and five that fed on uninfected leaves, co-inoculations involved five aphids that fed on PAV-inoculated culture leaves and five that fed on RPV-inoculated culture leaves (Appendix S1).

We destructively harvested the experimental plants at eight days post inoculation (DPI): 5, 8, 12, 16, 19, 22, 26, or 29 days. We cut the stems and leaves into small pieces using a sterilized blade, weighed them, stored about 60% of the tissue at −80°C for reverse transcription-quantitative polymerase chain reaction (RT-qPCR, see *Quantifying virus density*), and used the remainder to measure transmission (Fig. 1b). Unique combinations of nutrient treatments (*n* = 4), inoculation treatments (*n* = 3), and harvesting days (*n* = 8) resulted in 96 treatments. Each treatment was replicated twice in block one, once in blocks two through four, and zero to two times in block five, depending on losses in earlier blocks (Appendix S1: Table S2).

During blocks 1–4, approximately 40% of the tissue from the *Question 1* plants (i.e., “source plants”) was used to measure transmission to four “recipient plants” grown in each of the four nutrient treatments (*Question 2*, Fig. 1d). Source plant tissue was placed in glass tubes with 25 aphids for about two days. Then, five aphids, contained in a mesh cage affixed to the largest leaf of each recipient plant, fed for about four days (Appendix S1). The recipient plants were harvested 14 to 15 DPI and all of the stem and leaf tissue was stored at −80°C for RT-PCR and gel electrophoresis, which detects whether plants were infected with either virus (Appendix S1).

### Quantifying virus density

To quantify virus densities—number of viruses per milligram plant—in source plants, we first extracted the total RNA from ∼50 mg of thawed plant tissue (Appendix S1, Fig. 1c). We used one-step RT-qPCR to obtain the concentration of genomic RNA copies per volume of total RNA extract (Appendix S1). The RNA regions targeted for RT-qPCR, which are specific to PAV and RPV, encode coat proteins (Appendix S1: Table S3). We assumed that the genomic RNA copies measured by RT-qPCR approximated the number of virus particles in a sample and used the total amount of plant tissue extracted to estimate the concentration of viruses in 1 mg of plant tissue (Mackay et al. 2002, Lacroix et al. 2017). Virus densities that were large enough to be quantified by RT-qPCR were considered indicators of virus establishment. Plants with unintended infections (i.e., PAV detected in RPV-inoculated plants and RPV detected in PAV-inoculated plants, Appendix S1: Table S2) were excluded from analyses to generate conservative estimates. Such infections may have been caused by aphids that escaped inoculation cages or a small number of unintended infections in the virus culture leaves (Appendix S1).

### Statistical analysis

We performed all statistical analyses in R version 3.5.2 (R Core Team 2018), using the brms package (Bürkner 2017) to fit Bayesian linear regressions to data for each virus species. To evaluate virus establishment (*Question 1*), we fit a generalized linear regression with a Bernoulli response distribution (logit-link) to the proportion of plants that tested positive for infection based on RT-qPCR. To evaluate virus density in plants with infection (*Question 1*), we fit a normal linear regression to log-transformed virus density. In both cases, the predictor variables were a three-way interaction among the binary variables co-inoculation, N addition, and P addition (Appendix S2: Table S1). A first-order autocorrelation structure was used to account for correlations between virus density values on consecutive harvesting days. In the virus establishment models, harvesting day was included as a random intercept, which accounts for variation among harvesting days (autocorrelation structures were not compatible with this type of model). Experimental block was not included as a random intercept in either model because it explained minimal variation. We evaluated transmission (proportion of recipient plants infected, *Question 2*) using a generalized linear model with a Bernoulli response distribution (logit-link):

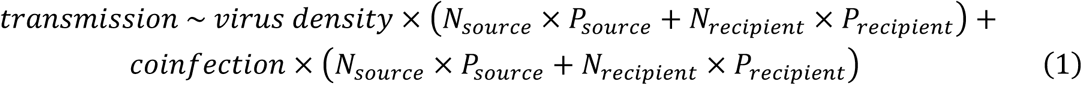

Subscripts indicate which plant the nutrient treatment was applied to and virus density was centered and scaled. Redundant main effects and interactions were omitted. Note that “co-infection” describes the status of the plant while “co-inoculation” describes the experimental treatment. Harvesting day and experimental block were included as crossed random intercepts. We used data from an experiment that measured PAV and RPV densities and transmission under similar conditions to inform some of the priors for the density and transmission models (Lacroix et al. 2017); uninformative priors were used otherwise (Appendix S2: Table S1). All models were run with three Markov chains, 6000 iterations each with a 1000 iteration warm-up. We evaluated model fit with r-hat values and visual comparisons of the observed data and simulated data from the posterior predictive distributions. We present the estimated effect sizes from models with informative priors, which were similar to models without informative priors (Appendix S2: Fig. S1). We report results as statistically significant if the 95% credible interval (CI; the interval that contains the most probable estimate values) omits the value representing “no effect” (i.e., zero for normal distribution or one for Bernoulli distribution).

### Mathematical model

To evaluate the effects of nutrition-mediated among-pathogen interactions on the spread of B/CYDVs in plant populations, we used our empirical results to parameterize a two-pathogen compartmental model (Seabloom et al. 2015). In the model, host plants are susceptible (*S*), infected with PAV (*I*_*P*_), infected with RPV (*I*_*R*_), or co-infected (*I*_*C*_):

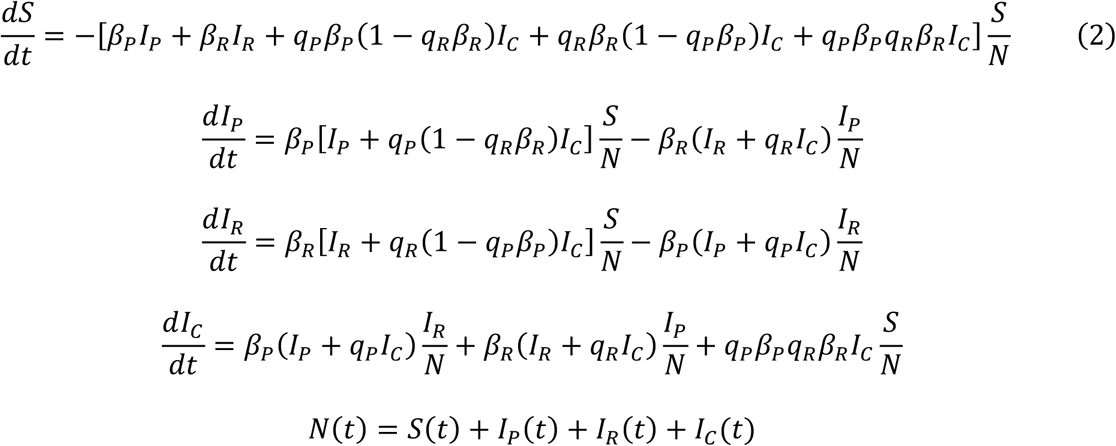

The terms *β*_*P*_ and *β*_*R*_ represent the probability of transmission from plants singly infected with PAV and RPV, respectively, given vector-assisted contact with another plant. Transmission from co-infected plants equals transmission from singly infected plants multiplied by a modifier (*q*_*P*_ or *q*_*R*_), which may represent positive (>1) or negative (<1) interactions (Appendix S3). We performed simulations of Eq. 2 over a single growing season (R version 3.5.2, R Core Team 2018) using the deSolve package (Soetaert et al. 2010). We compared simulations with both viruses present in the system to those with each virus alone. We repeated the simulations with three sets of parameter values estimated from Eq. 1 (Appendix S3: Table S1) that differ in the processes by which virus interactions can affect transmission: through changes in virus density, virus density–independent processes, and both types of processes (Appendix S3: Table S2). Parameter values for each nutrient treatment were used in separate simulations, restricting transmission to plants grown with the same nutrient treatment.

## Results

### Question 1: Does host nutrition affect among-pathogen interactions within hosts?

Nutrient addition and co-inoculation did not significantly affect PAV or RPV establishment (the proportion of plants infected; Appendix S2: Table S1). Co-inoculation had the strongest effects on PAV establishment when plants were grown with low nutrients (−29%, Fig. 2c) and RPV establishment when plants were grown with elevated N (+7%, Fig. 2d).

**Figure 2.**
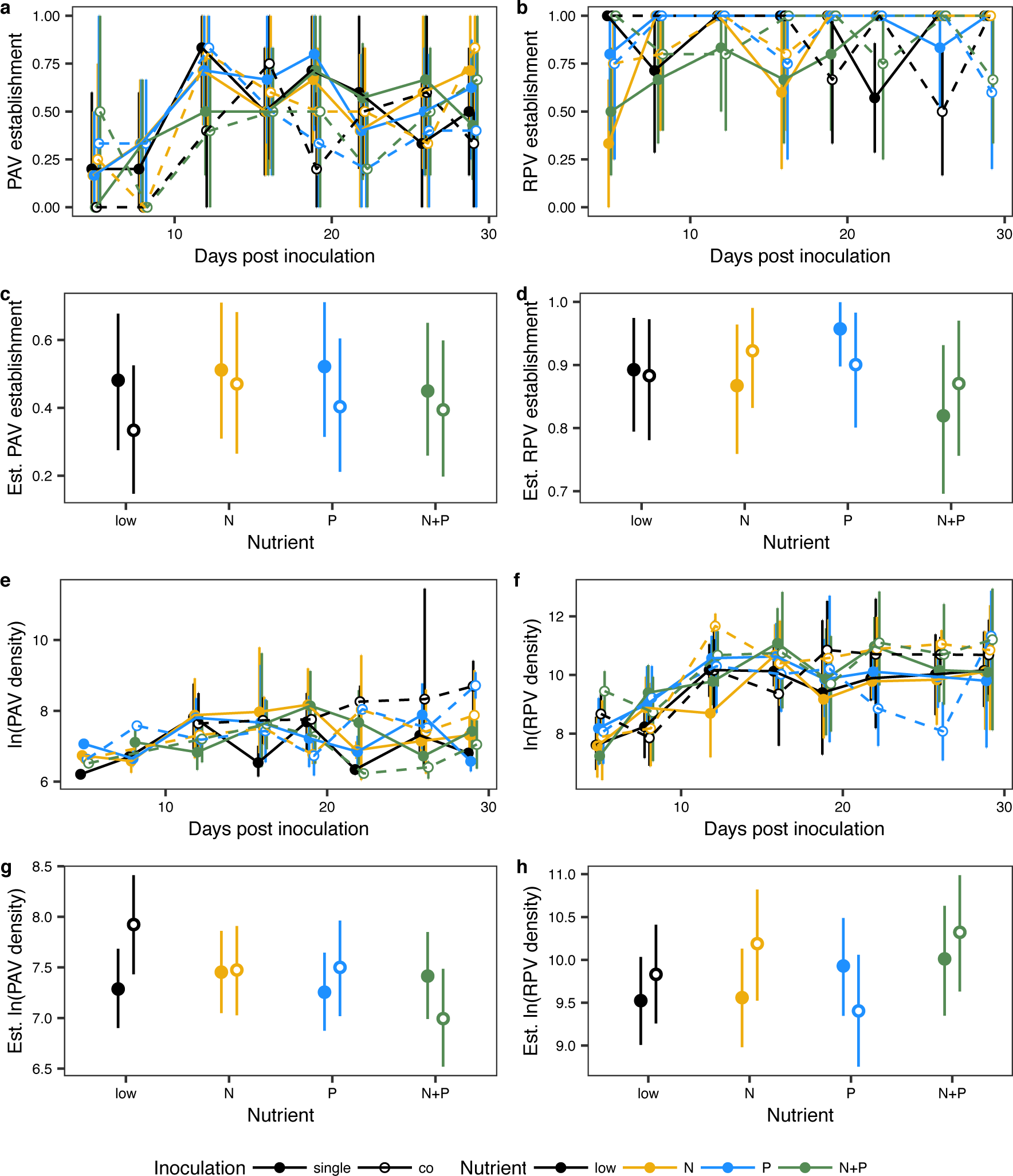
The effects of nutrients and co-inoculation on (a, b) establishment (proportion of source plants infected) and (e, f) log-transformed density (viruses per mg plant tissue) of (a, e) PAV and (b, f) RPV over time (mean ± 95% nonparametric bootstrap confidence intervals). Linear regression estimates of (c, g) PAV and (d, h) RPV (c, d) establishment and (g, h) log-transformed density (mean ± 95% credible intervals).

Nutrient addition and co-inoculation did not significantly affect RPV density (viruses per mg plant tissue; Table 1). Co-inoculation had the strongest effect on RPV density when plants were grown with elevated N (+105%, Fig. 2h). This positive effect was relatively consistent following the first two harvesting days (Fig. 2f). Co-inoculation significantly increased PAV density 98% when plants were grown with low nutrients (Fig. 2g, Table 1), which was more evident later in the course of infection (Fig. 2e).

**Table 1.** Model estimates and 95% credible intervals (CI) for statistical models of log-transformed virus density.

| Predictor | PAV |  | RPV |  |
| --- | --- | --- | --- | --- |
|  | Estimate | 95% CI | Estimate | 95% CI |
| co-inoculation | 0.64* | 0.06–1.22 | 0.31 | -0.36–0.97 |
| N addition (N) | 0.17 | -0.36–0.69 | 0.03 | -0.60–0.67 |
| P addition (P) | -0.03 | -0.54–0.47 | 0.40 | -0.29–1.10 |
| co-inoculation:N | -0.61 | -1.36–0.14 | 0.33 | -0.58–1.21 |
| co-inoculation:P | -0.39 | -1.12–0.35 | -0.83 | -1.82–0.14 |
| N:P | -0.01 | -0.72–0.70 | 0.05 | -0.91–0.98 |
| co-inoculation:N:P | -0.04 | -1.01–0.93 | 0.51 | -0.84–1.85 |

Note: Asterisk indicates estimate has 95% CI that do not include zero, which suggests that “no effect” is absent from the most probable estimate values.

### Question 2: Does host nutrition modify among-pathogen interactions to affect transmission?

Host nutrition modified the relationships between virus density and transmission (proportion of recipient plants infected; Table 2). RPV density significantly increased transmission when recipient plants were grown with elevated N and P (Fig. 3d). PAV displayed a similar trend (Fig. 3c). PAV density significantly decreased transmission when recipient plants were grown with elevated P (Fig. 3c); a trend also observed for RPV (Fig. 3d).

**Table 2.** Model estimates and 95% credible intervals (CI) for statistical models of virus transmission.

| Predictor | PAV |  | RPV |  |
| --- | --- | --- | --- | --- |
|  | Estimate | 95% CI | Estimate | 95% CI |
| density | 0.82 | 0.52–1.28 | 1.30* | 1.03–1.63 |
| co-infection | 0.68 | 0.14–2.99 | 0.84 | 0.46–1.56 |
| N addition to source ($N_{\text{source}}$) | 1.72 | 0.72–4.15 | 0.92 | 0.54–1.60 |
| P addition to source ($P_{\text{source}}$) | 0.83 | 0.35–1.91 | 1.60 | 0.85–3.04 |
| N addition to recipient ($N_{\text{recipient}}$) | 0.75 | 0.20–2.83 | 2.81* | 1.25–6.69 |
| P addition to recipient ($P_{\text{recipient}}$) | 0.18* | 0.05–0.57 | 1.33 | 0.64–2.86 |
| $N_{\text{source}}:P_{\text{source}}$ | 0.79 | 0.24–2.53 | 0.66 | 0.29–1.49 |
| $N_{\text{recipient}}:P_{\text{recipient}}$ | 8.88* | 1.35–64.55 | 0.9 | 0.26–3.12 |
| density:$N_{\text{source}}$ | 2.01 | 0.95–5.13 | 0.77 | 0.22–2.56 |
| density:$P_{\text{source}}$ | 2.72 | 0.84–10.55 | 1.19 | 0.70–2.26 |
| density:$N_{\text{recipient}}$ | 1.36 | 0.50–4.95 | 0.78 | 0.47–1.38 |
| density:$P_{\text{recipient}}$ | 0.44* | 0.17–0.95 | 0.68 | 0.41–1.10 |
| co-infection:$N_{\text{source}}$ | 0.22* | 0.05–0.93 | 1.87 | 0.90–3.85 |
| co-infection:$P_{\text{source}}$ | 0.63 | 0.14–2.72 | 0.82 | 0.37–1.79 |
| co-infection:$N_{\text{recipient}}$ | 1.74 | 0.35–8.54 | 0.87 | 0.29–2.60 |
| co-infection:$P_{\text{recipient}}$ | 8.60* | 2.00–41.27 | 0.77 | 0.29–1.99 |
| density:$N_{\text{source}}:P_{\text{source}}$ | 0.13 | 0.01–1.50 | 0.63 | 0.15–2.49 |
| density:$N_{\text{recipient}}:P_{\text{recipient}}$ | 7.24 | 0.88–110.44 | 4.27* | 1.57–15.61 |
| co-infection:$N_{\text{source}}:P_{\text{source}}$ | 1.71 | 0.27–11.02 | 1.21 | 0.41–3.66 |
| co-infection:$N_{\text{recipient}}:P_{\text{recipient}}$ | 0.17 | 0.02–1.58 | 0.81 | 0.16–4.16 |

Note: Asterisk indicates estimate (odds ratios) has 95% CI that do not include one, which suggests that “no effect” is absent from the most probable estimate values.

**Figure 3.**
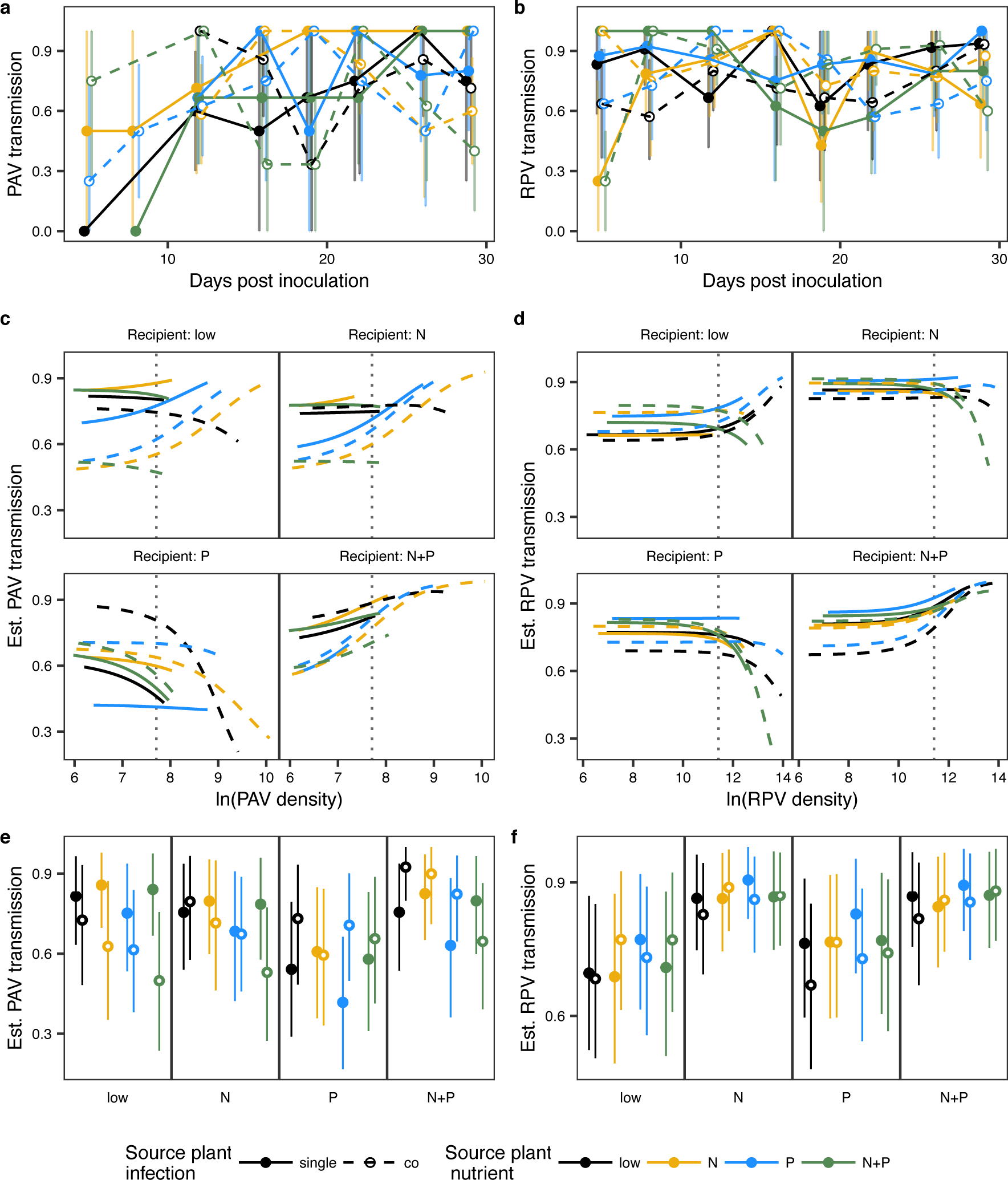
The effects of source plant nutrition and infection status on (a) PAV and (b) RPV transmission (proportion of recipient plants infected) over time, averaged over recipient plant nutrient treatments (mean ± 95% nonparametric bootstrap confidence intervals; see Appendix S2: Fig. S2 for averages over source plants). Regression relationships between transmission and log-transformed virus density for (c) PAV (d) RPV. Vertical lines indicate overall average log-transformed density for each virus. Regression estimates of (e) PAV and (f) RPV transmission (mean ± 95% credible intervals) were taken at the average virus density for each treatment.

Consistent with the results for PAV density (Fig. 2g), co-infection significantly increased PAV transmission 43% when source plants were grown with low nutrients and recipient plants were grown with elevated P (Fig. 3e, Table 2). However, PAV density reduced transmission under these conditions (Fig. 3c) and co-infection increased PAV transmission 93% when density was held constant (i.e., the difference in transmission at the vertical dotted line on Fig. 3c). Co-infection significantly reduced PAV transmission 26% from plants grown with elevated N to plants grown with low nutrients, with stronger effects when density was held constant (−38%, Fig. 3e). Nitrogen addition to recipient plants significantly increased RPV transmission 26% (source plants grown with low nutrients; Fig. 3f). Co-infection did not significantly affect RPV transmission, increasing it the most when source plants were grown with elevated N and recipient plants were grown with low nutrients (14%, Fig. 3f). This result is consistent with the positive effect of co-inoculation on RPV density (Fig. 2h), but co-infection still increased transmission 14% when density was held constant.

### Question 3: Can nutrition-mediated among-pathogen interactions affect disease spread?

Simulations from the mathematical model (Eq. 2) suggest that RPV can increase PAV infection prevalence in plant populations grown with elevated P and decrease PAV prevalence with the addition of N or both nutrients (Fig. 4a). These effects are driven by among-pathogen interactions that do not act on transmission through changes in virus density (i.e., density-independent, Fig. 4c). In fact, interactions with RPV that alter PAV density increase PAV infection prevalence with N addition (Fig. 4b). Simulations suggest that PAV can increase RPV infection prevalence with N addition and decrease RPV infection prevalence when plants are grown with low nutrients (Fig. 4d). Again, these effects are driven by virus density–independent processes (Fig. 4f) and changes in density due to among-pathogen interactions have some opposite effects (Fig. 4e). The predicted effects begin about midway through the growing season and later decline as all plants in the population become infected (Appendix S3: Fig. S1).

**Figure 4.**
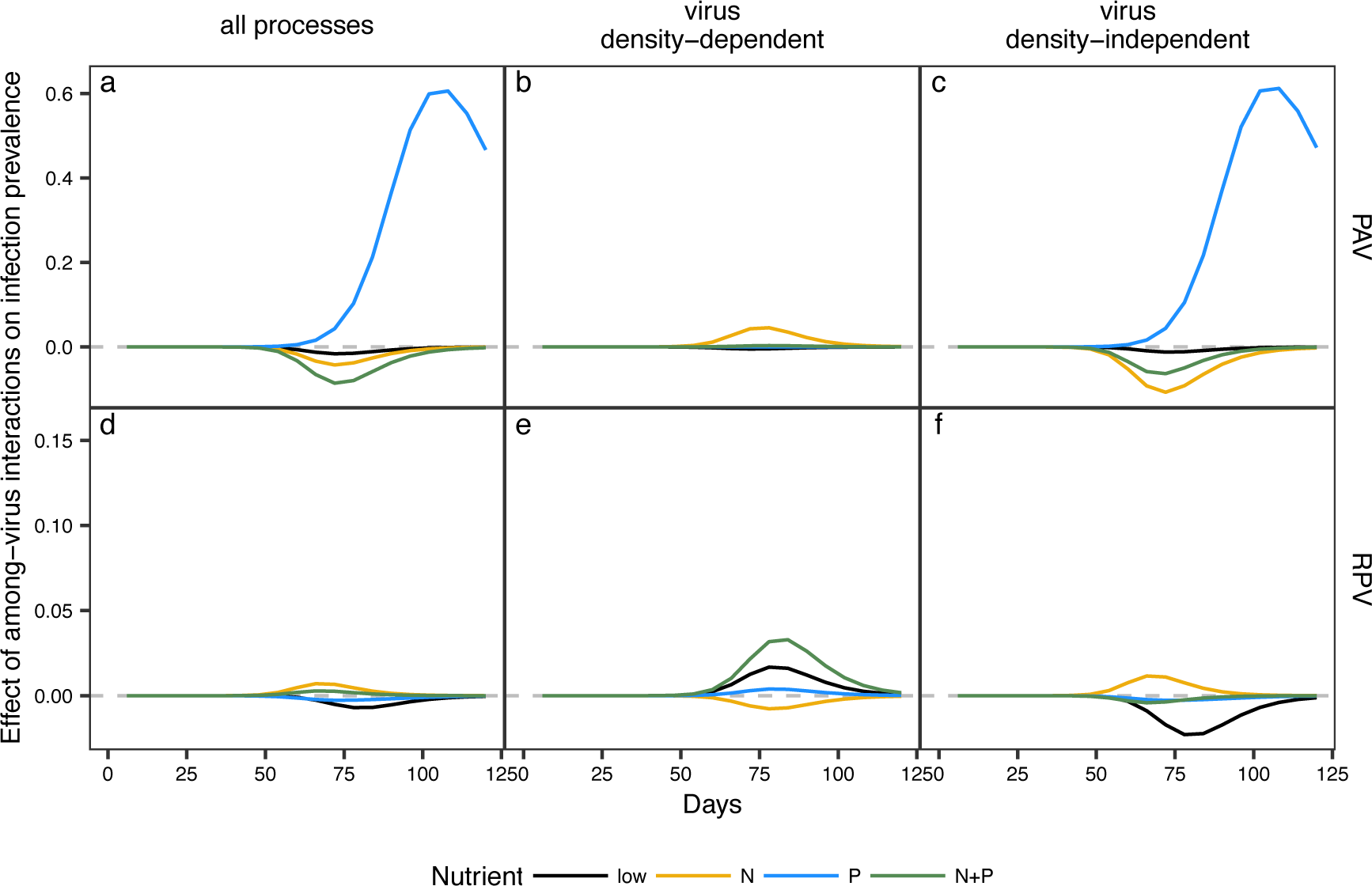
The predicted change in prevalence of (a–c) PAV and (d–f) RPV due to the presence of the other virus in simulated plant populations grown with low nutrients, N addition, P addition, or both nutrients. Initial host population sizes were *I*_*P*_ = 1 (when PAV was present), *I*_*R*_ = 1 (when RPV was present), *I*_*C*_ = 0, *N* = 4,000. Parameter values represent transmission that (b, e) depends on virus density, (c, f) is independent of virus density, or (a, d) both.

## Discussion

The results from this experiment are consistent with findings from previous studies across host taxa: host nutrition can mediate within-host interactions among pathogens (Lacroix et al. 2014, Lange et al. 2014, Budischak et al. 2015, Wale et al. 2017). We built upon this work to demonstrate that host nutrition and among-pathogen interactions can alter transmission, and potentially disease spread. In particular, plant viruses can promote one another’s within-host densities under specific host nutrition conditions (*Question 1*). The viruses had positive, negative, and neutral effects on one another’s transmission, which also varied with host nutrition (*Question 2*). A mathematical model parameterized with these experimental results suggests that interactions between viruses that alter transmission directly—as opposed to indirectly through changes in density—will affect disease spread in a population (*Question 3*).

### The effects of host nutrition on among-pathogen interactions within the host

Changes in host nutrition shifted among-pathogen interactions from neutral to positive. With low nutrients, co-inoculation slowed the establishment of PAV, but ultimately promoted PAV density. In contrast, co-inoculation had limited effects on PAV with elevated N and P supplies. Co-inoculation increased RPV density with N addition. These results suggest that N and P supplied to grasses through fertilization, atmospheric deposition, and other processes, may alter the strength of interactions between viruses co-occurring within hosts. We used previous studies on nutrition-mediated PAV and RPV interactions to inform the priors of our statistical models. In the previous studies, elevated N alleviated a negative effect of co-inoculation on RPV establishment, but host nutrition did not mediate the effects of co-inoculation on virus density (Lacroix et al. 2014, 2017). We also found that N addition led to a more positive effect of co-inoculation on RPV. Our result that co-inoculation increased PAV density with low nutrient supply provided a novel insight into our understanding of among-pathogen interactions.

Our work is consistent with previous studies that have shown that B/CYDVs, like other plant viruses, can have positive effects on one another. Although it is not yet known how PAV and RPV apparently facilitate each other, there are at least two potential explanations. Plant viruses can use proteins produced by other virus species, which may facilitate transmission and movement through the plant (Wen and Lister 1991, Moreno and López-Moya 2020). Different plant viruses also can interfere with host immunity using distinct mechanisms (Liu et al. 2012). Complementarity in host immunosuppression may increase virus density (Moreno and López-Moya 2020). Both mechanisms—sharing resources and complementary immunosuppression— may be mediated by host nutrition (Smith et al. 2005, Budischak et al. 2015).

### The effects of nutrition-mediated among-pathogen interactions on transmission

Host nutrition mediated the size and direction of the effects of among-pathogen interactions on transmission. Although the positive effects of co-infection on transmission occurred under the same nutrient treatments as positive effects of co-inoculation on density, higher virus density did not explain increased transmission. In fact, the relationship between virus density and transmission was variable for both viruses. This results is similar to a field experiment manipulating plant fungal infection in which plants with more infected leaves did not consistently produce more fungal spores (Susi et al. 2015b). Also, the density-transmission relationship depended upon host nutrition, a result that is parallel to nutrition effects on aphid endosymbionts (Wilkinson et al. 2007), suggesting that these interdependencies may be general.

A range of factors other than within-host pathogen density can affect transmission (McCallum et al. 2017). B/CYDV transmission depends on virus-vector and host-vector interactions (Rochow et al. 1983, Gray et al. 1991, Wen and Lister 1991). Highly relevant to our results are the findings that plant nutrient content and infection status affect aphid feeding preferences (Srinivasan and Alvarez 2007, Nowak and Komor 2010) and that the time aphids spend feeding on plants can affect transmission (Gray et al. 1991). While we partially controlled for aphid preference by placing aphids in cages, we do not know how long aphids fed on each plant. Thus, observed changes in transmission due to host nutrition and co-infection may have arisen through variation in aphid feeding times. In this case, the presence of one pathogen can affect the fitness and prevalence of the other (i.e., an “among-pathogen interaction”), despite the absence of relevant changes in within-host density.

### The implications of host nutrition-mediated among-pathogen interactions for disease spread

A mathematical model parameterized with empirical transmission values demonstrated that nutrition-mediated among-pathogen interactions may affect infection prevalence in plant populations and highlighted the importance of virus density–independent processes in transmission. The result that changes in within-host virus density due to nutrition-mediated among-pathogen interactions are unlikely to affect infectious disease dynamics in host populations is consistent with research in animal populations demonstrating that nutrients can influence infection prevalence through transmission processes that are independent of within-host pathogen density, such as contact between susceptible and infectious hosts (Becker et al. 2015). Evaluating the relative effects of within-host dynamics and other transmission-related processes on infectious disease dynamics is an important goal of disease ecology, especially considering the complexity that within-host dynamics can add to empirical and theoretical studies (Mideo et al. 2008, Handel and Rohani 2015, Susi et al. 2015b).

Nonetheless, some of the predictions of the model were not apparently consistent with previous work. In two separate field experiments, P addition, but not N addition, increased PAV prevalence, and in one experiment, neither nutrient affected RPV prevalence (Seabloom et al. 2013, Borer et al. 2014). Our model predicted that P addition would reduce PAV prevalence, despite positive effects of co-infection under this condition. Both N and P were predicted to increase RPV prevalence. The effects of co-infection and host nutrition on aphid preference, aphid population growth, and other factors affecting transmission that were not measured in this experiment may explain the gap between model predictions and field experiment results. Experiments examining such processes (e.g., Srinivasan and Alvarez 2007, Nowak and Komor 2010) could be paired with more detailed models (e.g., Strauss et al. 2019) to further explore the implications of nutrition-mediated among-pathogen interactions for infectious disease dynamics.

### Limitations of this study

We observed high uncertainty around estimates of establishment, pathogen density, and transmission, which may result from variation in host-pathogen interactions across individuals (de Roode et al. 2004). This variation could be amplified in our dataset if it is more apparent when viruses reach higher densities: the lower detection threshold of our RT-qPCR protocol (about 150 viruses per mg plant) limited our ability to accurately quantify samples with low virus densities, leading to their removal from density and transmission analyses. In addition, we conducted simultaneous inoculations of PAV and RPV, but the sequence and timing of inoculations can affect the outcome of pathogen interactions (Clay et al. 2018). Host nutrition may have different effects on pathogen interactions depending on inoculation sequence and timing. Nonetheless, our results do empirically demonstrate that a host’s nutritional environment can alter among-pathogen interactions, transmission, and disease spread.

### Conclusions

This study demonstrates that host nutrition may alter infectious disease dynamics through among-pathogen interactions. Influential nutrition-mediated among-pathogen interactions manifested as changes in transmission that were independent of within-host pathogen density. Therefore, the development of a more comprehensive, predictive framework for the role of co-infection in disease transmission and infectious dynamics would benefit from investigations of host nutrition effects on virus-vector and host-vector interactions (Rochow et al. 1983, Nowak and Komor 2010). Co-infection of hosts is common in natural systems (Tollenaere et al. 2016), where host nutrition is altered by intentional and unintentional nutrient inputs (Smith et al. 2005). Overall, the results from this study suggest that nutrient inputs into terrestrial plant systems are likely to affect interactions between co-occurring viruses, leading to shifts in disease spread.

## Acknowledgements

We are grateful to Christelle Lacroix, Melissa Rudeen, Anita Krause, Nicholas Cupery, Casey Easterday, Kurra Renner, Luc Robichaud, and Alexis Rogers for help with the experiment and to multiple anonymous reviewers for their comments on earlier drafts. AEK was supported by an NSF IGERT graduate fellowship at the University of Minnesota (DGE-0653827) and an NSF Graduate Research Fellowship (base award number 006595) and ETB and EWS received support from the NSF program in Ecology and Evolution of Infectious Diseases (grant DEB-1015805). AEK, ETB, and EWS designed the experiment, AEK, ENB, and TCP performed the experiment and analyses, AEK wrote the first draft, and all authors contributed to revisions.

## Data availability

Data and code are publicly available on the Environmental Data Initiative Data Portal: https://doi.org/10.6073/pasta/01e7bf593676a942f262623710acba13

