## Appendix S1 for "Host nutrition mediates interactions between plant viruses, altering transmission and predicted disease spread"

**Appendix S1: Extended methods for experimental design and data collection**
"Host nutrition mediates interactions between plant viruses, altering transmission and predicted disease spread"
Amy E. Kendig, Elizabeth T. Borer, Emily N. Boak, Tashina C. Picard, and Eric W. Seabloom
*Ecology*

*Inoculating plants with viruses*

*Rhopalosiphum padi* were starved in sealed glass tubes in groups of twenty for up to four hours to help ensure feeding on target plants. We then added 39.0 mg ± 10.3 mg of uninfected, PAV-inoculated, or RPV-inoculated leaves from virus cultures to the tubes. After 44 to 48 hours, we pooled the aphids in large plastic containers according to their inoculation type and added them to 2.5 x 8.5 cm, 118 µm polyester mesh cages (Sefar America Inc., Kansas City, MO, USA) affixed to the largest leaf on each plant. The cages were secured to the leaves with Parafilm® (Bemis Company, Inc., Neenah, WI, USA) and two bobby pins on each end. The aphids were allowed to feed for approximately four days: 98, 100, 101, 120.25, and 98.25 hours for each of the experimental rounds. While feeding for the fourth round lasted one day longer, plants were still harvested the same number of days from the initiation of aphid feeding as the other rounds. When the feeding time was finished, the aphids were manually crushed, the cages were removed, and the leaves were wiped with a gloved hand to remove remaining aphids. Aphids occasionally survived, presumably because they were small enough to escape the cage or avoid crushing (e.g., offspring of the aphids inserted in the cage, which were often observed). We therefore periodically checked plants and manually removed aphids throughout the experiment. We also placed yellow sticky traps in the growth chamber to catch winged aphids or other insects. Few insects were caught with these traps and none, besides aphids, were observed on the experimental plants. The first day the aphids were placed on plants is designated "day 0" in the calculation of days post inoculation (DPI).

Transmission from source plants to recipient plants occurred at each harvesting day during blocks 1-4 and followed the same inoculation methods described above with the following exceptions. For the aphids to acquire viruses from the source plants, we placed 25 aphids in sealed tubes with 40% of the leaf and stem tissue. During block 1, there was a shortage of aphids for the second and third harvesting days, so we used 13 to 25 aphids per tube and scaled down the plant tissue accordingly. Aphids fed on source plant tissue for 38 to 55 hours. Five aphids were used to inoculate each recipient plant. We aimed to inoculate four recipient plants for each source plant, one grown in each nutrient treatment. However, inoculations occasionally failed because the leaf with aphids affixed to it broke off. Aphids fed on recipient plants for 86 to 103 hours.

The virus culture leaves used to inoculation source plants could not be tested for infection until after inoculation of the experimental plants. We used RNA extraction, reverse transcription, polymerase chain reaction (RT-PCR), and gel electrophoresis to assess their infection status and found that 93% of PAV-inoculated culture leaves tested positive for PAV infection, 88% of RPV-inoculated culture leaves tested positive for RPV infection, and 10% of all culture leaves tested positive for PAV or RPV when those infections were not intended. The virus densities in the culture leaves were not quantified, but 90% of the unintended infections had gel electrophoresis bands that were lighter than the positive controls, suggesting lower virus density, in contrast to 9% and 12% of intentional PAV and RPV infections, respectively.

*Detecting viruses with RT-PCR*

To detect the presence of PAV and RPV in plants, we used RNA extraction, reverse transcription (RT), polymerase chain reaction (PCR), and gel electrophoresis. Between each step, samples were stored at -20°C. The methods used for RNA extraction apply to tissue used for virus detection (this section) and virus density quantification. The combined stem and leaf tissue for each plant was cut into small pieces with a sterilized blade. Fifty milligrams of haphazardly selected segments were ground up with 1 mL TRIzol™ Reagent (Invitrogen™, Thermo Fisher Scientific, Waltham, MA, USA) in a bead beater. The TRIzol supernatant was separated from the remaining tissue components through centrifugation. Chloroform was added and the samples were centrifuged to separate the RNA from the DNA and proteins. The RNA was then precipitated from the aqueous layer using isopropanol and centrifugation. The RNA was washed with 75% ethanol and dissolved in 50 µl of RNase-free water (Qiagen).

The total RNA concentration was measured using a nanodrop spectrophotometer (Thermo Fisher Scientific) and used to determine the volume of RNA to use in RT. Dissolved RNA was combined with random hexamers, which are generic primers that lead to the reverse transcription of all RNA in the sample, including plant RNA. This mixture was denatured by heating it up to 70°C. Then, 5X ImProm-II reaction buffer, 25 mMMgCl$_2$, 10 mM dNTPs, 20 U RNasin, Improm-II Reverse Transcriptase (all from Promega™), and RNase-free water (Qiagen) were added to the samples. The mixture was put in the thermal cycler and heated to 25°C for 5 minutes for low temperature RT, which prevents the random hexamers from separating from the RNA. Then, optimum temperature RT took place at 45°C for 60 minutes. This was followed by deactivation of the reverse transcriptase at 70°C for 15 minutes.

The cDNA product from RT was combined with 10X PCR Buffer (Qiagen), 25 mM MgCl$_2$ (Promega™), 10 mM dNTPs (Promega™), 1 U HotStart Taq (Qiagen), 10 µM forward and reverse primers for PAV (PAVR1, ATTGTGAAGGAATTAATGTA; PAVL1, AGAGG-AGGGGCAAATCCTGT), 5 µM forward and reverse primers for RPV (RPVR2 CTGCGTTCTGACAGCAGG, RPV L ATGTTGTACCGC- TTGATCCAC), and water (Deb and Anderson 2008, Lacroix et al. 2014). In the thermal cycler, the Taq polymerase was activated with a 15-minute period of 95°C. The cDNA was then amplified by six cycles of denaturation of DNA at 95°C for 30 seconds, annealing of primers to ssDNA at 66°C for 30 seconds (with the temperature decreasing by 1°C for each cycle), and extension of primers at 72°C for 1 minute. With the annealing step lasting for 1 minute, the same denaturation, annealing, and extension stages were repeated 30 more times to amplify the DNA. There was a final extension step of 72°C for 10 minutes.

Five microliters of each PCR product were combined with 2 µL of 6X loading dye (Genesee Scientific). The samples were loaded into a fine-resolution SybrSafe (Invitrogen, Thermo Fisher Scientific)-stained 2% (w/v) Agarose-1000 (Invitrogen, Thermo Fisher Scientific) gel. After approximately 25 minutes, the gel was analyzed with a UV-light EZ doc system (Bio-Rad Laboratories). The fragment size (RPV, 447 bp; PAV, 298 bp) was checked relative to a 100 bp DNA ladder (Apex BioResearch Products). We compared visible bands to positive controls and visually assessed whether the band was lighter or darker than the control. We explored the implications of using only the darker bands as evidence of infection for the analysis of the transmission dataset and found qualitatively similar results to when all visible bands were taken as evidence for infection (see archived code). Therefore, we assumed that any visible band is evidence of infection in all relevant analyses (i.e., assessing unintended infections and evaluating treatment effects on transmission).

*Quantifying virus density with RT-qPCR*

To estimate the density of each virus species within the plants, we used RNA extraction and one-step reverse transcription-quantitative polymerase chain reaction (RT-qPCR). Fifty microliters of RNA extract were produced for each sample following the methods described above and stored at -80$^{o}$C.

To prepare RT-qPCR standards for each virus, we extracted RNA from *A. sativa* tissue with known infections, performed an RT-PCR (Appendix S3), and converted the product to RNA (MEGAscript® T7 Kit, Applied Biosystems®), which was diluted to a series of $10^7$ to $10^3$ copies of viral RNA. We also designed forward and reverse qPCR primers and TaqMan® probes for each virus that target regions encoding the coat protein (Appendix S1: Table S3).

To prepare the RT-qPCR reactions, seven microliters of RNA (or water, for the negative controls) were combined with 0.25 μM TaqMan® probes, 0.3 μM of forward and reverse primers for PAV or RPV (Appendix S1: Table S3), 1X buffer, and 1X enzyme mixture (RNA-to-$C_T$™ 1-Step Kit, Applied Biosystems®, Thermo Fisher Scientific). The reaction was kept at 48$^{o}$C for 30 minutes for the RT, 95$^{o}$C for 10 minutes for activation of Taq polymerase, and 50 cycles of 95$^{o}$C for 25 seconds and 60$^{o}$C for one minute were repeated for denaturation, annealing, and extension. Co-inoculated source plants and many of the RPV-inoculated source plants were analyzed twice–once for PAV and once for RPV. Most PAV-inoculated source plants were only analyzed for PAV, and RT-PCR was used to check for the presence of RPV.

Each sample was replicated three times in a 96-well plate, which was analyzed with the Applied Biosystems® StepOnePlus™ Real-time PCR System. The detection threshold was set automatically by the system. Standard curves were used to calculate efficiency, with 100% efficiency indicating that DNA strands approximately doubled in quantity with each PCR cycle (Bustin et al. 2009). We excluded standards from analysis if they had a lower density than any detected negative controls (indicating a potential false positive). Samples in runs with standard curves outside the target efficiency range (85% - 115%) were re-analyzed when possible but excluded from analyses otherwise. We estimated the quantity of RNA within a well using the detection threshold and standard curve. To estimate the density of PAV or RPV in a sample (number per mg plant), we took the mean of the concentration (genomic copies/μl) in the technical replicates (wells), multiplied by 50 μl (the volume of the RNA extract) to estimate the copies in the RNA extract, and then divided by the mass of plant tissue used for the RNA extraction.

Some of the samples were quantified by RT-qPCR, but they fell below the lowest density on the standard curve. We attempted to increase the range of the standard curve to include samples with $10^2$ copies/well, but standards prepared at this density were not consistently detected. In fact, some of the RT-qPCR runs did not even detect standards prepared with $10^3$ copies/wel1. When possible, we re-analyzed samples that fell below the standard curve, especially when the standards with $10^3$ copies/wel1 were not detected. When calculating the mean of well replicates, we removed replicates that were detected, but outside of the standard curve. Non-detected replicates were included in the mean as zero density and non-detected samples (i.e. all replicates were not detected) were included in the analysis of infection rate, but not virus density. Only source plants with quantified virus densities corresponding to their inoculation treatment were used in the analysis of transmission.

**Appendix S1: Table S1.** Modified Hoagland solution recipe used to create nutrient treatments

| Compound | Molar Mass | Concentration (μM) |
|---|---|---|
| $K_2SO_4$ | 174.25 | 1250 |
| $MgSO_4{\cdot}7H_2O$ | 246.48 | 1000 |
| $KH_2PO_4$ | 136.09 | 1 (50 for elevated P) |
| $CaSO_4{\cdot}2H_2O$ | 172.17 | 2000 |
| $NH_4NO_3$ | 80.04 | 7.5 (375 for elevated N) |
| KCl | 74.56 | 25 |
| $H_3BO_3$ | 61.83 | 12.5 |
| $MnSO_4{\cdot}H_2O$ | 169.02 | 1 |
| $ZnSO_4{\cdot}7H_2O$ | 278.56 | 1 |
| $CuSO_4{\cdot}5H_2O$ | 249.69 | 0.25 |
| $H_2MoO_4{\cdot}(H_2O)$ | 161.95 | 0.25 |
| NaFeEDDHA (6% Fe) | 434.80 | 10 |

Note: The low and high concentrations of $KH_2PO_4$ and $NH_4NO_3$ are 0.2% and 10% of half-strength Hoagland solution, respectively (Hoagland and Arnon 1938, Seabloom et al. 2011, Lacroix et al. 2014, 2017).

**Appendix S1: Table S2.** Sample sizes

| Plant | Nutrient | Inoculation | PAV | Unintended RPV | RPV | Unintended PAV |
|---|---|---|---|---|---|---|
| source | low | single | 47 (22) | 2 | 48 (38) | 2 |
| source | N | single | 48 (23) | 2 | 47 (35) | 4 |
| source | P | single | 49 (25) | 1 | 48 (43) | 1 |
| source | N + P | single | 49 (23) | 1 | 49 (37) | 2 |
| source | low | co | 42 (14) | - | 42 (37) | - |
| source | N | co | 41 (21) | - | 41 (34) | - |
| source | P | co | 42 (18) | - | 42 (35) | - |
| source | N + P | co | 41 (17) | - | 41 (34) | - |
| recipient | low → low | single | 17 (08) | 5 | 26 (20) | 1 |
| recipient | low → N | single | 16 (09) | 3 | 27 (21) | 1 |
| recipient | low → P | single | 11 (07) | 2 | 25 (21) | 0 |
| recipient | low → N + P | single | 15 (08) | 4 | 24 (19) | 1 |
| recipient | N → low | single | 10 (07) | 1 | 21 (12) | 3 |
| recipient | N → N | single | 15 (09) | 2 | 26 (18) | 2 |
| recipient | N → P | single | 14 (09) | 1 | 25 (16) | 2 |
| recipient | N → N + P | single | 17 (08) | 4 | 25 (16) | 3 |
| recipient | P → low | single | 20 (12) | 3 | 29 (21) | 3 |
| recipient | P → N | single | 17 (08) | 4 | 26 (20) | 2 |
| recipient | P → P | single | 19 (11) | 4 | 28 (20) | 3 |
| recipient | P → N + P | single | 14 (05) | 4 | 28 (18) | 4 |
| recipient | N + P → low | single | 17 (10) | 4 | 24 (18) | 2 |
| recipient | N + P → N | single | 15 (09) | 4 | 23 (19) | 1 |
| recipient | N + P → P | single | 16 (09) | 4 | 26 (20) | 1 |
| recipient | N + P → N + P | single | 17 (10) | 3 | 29 (21) | 0 |
| recipient | low → low | co | 28 (10) | - | 28 (26) | - |
| recipient | low → N | co | 28 (09) | - | 28 (25) | - |
| recipient | low → P | co | 25 (10) | - | 25 (23) | - |
| recipient | low → N + P | co | 29 (10) | - | 29 (27) | - |
| recipient | N → low | co | 22 (13) | - | 22 (15) | - |
| recipient | N → N | co | 29 (14) | - | 29 (23) | - |
| recipient | N → P | co | 24 (12) | - | 24 (17) | - |
| recipient | N → N + P | co | 27 (13) | - | 27 (21) | - |
| recipient | P → low | co | 30 (15) | - | 30 (25) | - |
| recipient | P → N | co | 27 (14) | - | 27 (23) | - |
| recipient | P → P | co | 30 (15) | - | 30 (24) | - |
| recipient | P → N + P | co | 27 (14) | - | 27 (21) | - |
| recipient | N + P → low | co | 26 (08) | - | 26 (20) | - |
| recipient | N + P → N | co | 26 (12) | - | 26 (21) | - |
| recipient | N + P → P | co | 24 (10) | - | 24 (20) | - |
| recipient | N + P → N + P | co | 30 (12) | - | 30 (24) | - |

Note: PAV and RPV indicate sample sizes for the experiment and for virus density measurements (in parentheses), both pooled across harvesting days and experimental blocks. Unintended RPV and Unintended PAV are the number of unintended infections (RPV infections of PAV-inoculated plants, PAV infections of RPV-inoculated plants, respectively). Virus density sample sizes are smaller than experiment sample sizes because the plants were not infected, the virus density could not be reliably quantified, or the plants had unintended infections.

**Appendix S1: Table S3.** Sequences for RT-qPCR primers and probes

| Virus | NCBI accession | Sequence type | Sequence (5' to 3') |
|---|---|---|---|
| RPV | L25299 (Vincent et al. 1991) | Forward primer | GAGGTTAGCGAGGAGTTAGAATTC |
| | | Reverse primer | AACTACCTCAGAGTTGCCACATTC |
| | | Probe | VICACATCTTCAAGACTCCTAACCTCGCCATMGBNFQ |
| PAV | D11032 (Ueng et al. 1992) | Forward primer | TGGTCGCCCAAAAATCTAAAAC |
| | | Reverse primer | GGAGTAAGGCTCGCAGTAAATTGCCGCATAAACAC |
| | | Probe | FAMGGTGACCGAGGCTTGGACCGACTTCTTTMGBNFQ |

Notes: Primers and probes first reported in Lacroix et al. (2017) and adapted from Deb and Anderson (2008). We added a T7 promoter to the 5' end of the reverse primers (5′-TAATACGACTCACTATAGGGAGA-3′) for producing standards with RT-PCR prior to transcription (Lacroix et al. 2017).

**Appendix S1 literature cited**

Bustin, S., V. Benes, J. A. Garson, J. Hellemans, J. Huggett, M. Kubista, R. Mueller, T. Nolan, M. W. Pfaffl, G. L. Shipley, J. Vandesompele, and C. T. Wittwer. 2009. The MIQE guidelines: Minimum information for publication of quantitative real-time PCR experiments. Clinical Chemistry 55:611–622.

Deb, M., and J. M. Anderson. 2008. Development of a multiplexed PCR detection method for Barley and Cereal yellow dwarf viruses, Wheat spindle streak virus, Wheat streak mosaic virus and Soil-borne wheat mosaic virus. Journal of Virological Methods 148:17–24.

Hoagland, D. R., and D. I. Arnon. 1938. The water culture method for growing plants without soil. California Agricultural Experiment Station Circular 347:32.

Lacroix, C., E. W. Seabloom, and E. T. Borer. 2014. Environmental nutrient supply alters prevalence and weakens competitive interactions among coinfecting viruses. The New Phytologist 204:424–433.

Lacroix, C., E. W. Seabloom, and E. T. Borer. 2017. Environmental nutrient supply directly alters plant traits but indirectly determines virus growth rate. Frontiers in Microbiology 8:2116.

Seabloom, E. W., C. D. Benfield, E. T. Borer, A. G. Stanley, T. N. Kaye, and P. W. Dunwiddie. 2011. Provenance, life span, and phylogeny do not affect grass species' responses to nitrogen and phosphorus. Ecological Applications 21:2129–2142.

Ueng, P. P., J. R. Vincent, E. E. Kawata, C. H. Lei, R. M. Lister, and B. A. Larkins. 1992. Nucleotide sequence analysis of the genomes of the MAV-PS1 and P-PAV isolates of barley yellow dwarf virus. Journal of General Virology 73:487–492.

Vincent, J. R., R. M. Lister, and B. A. Larkins. 1991. Nucleotide sequence analysis and genomic organization of the NY-RPV isolate of barley yellow dwarf virus. The Journal of General Virology 72:2347–2355.
