## Appendix S2 for "Host nutrition mediates interactions between plant viruses, altering transmission and predicted disease spread"

**Appendix S2: Extended methods and results for statistical analyses**
"Host nutrition mediates interactions between plant viruses, altering transmission and predicted disease spread"
Amy E. Kendig, Elizabeth T. Borer, Emily N. Boak, Tashina C. Picard, and Eric W. Seabloom
*Ecology*

**Appendix S2: Table S1.** Statistical models

| Response | Predictor | Estimate | CI (95%) | Prior dist. | Hyperparameters |
|---|---|---|---|---|---|
| PAV establishment | intercept | 0.92 | 0.39–2.16 | normal | $\mu = 0$, $\sigma^2 = 10$ |
| PAV establishment | co-inoculation | 0.53 | 0.21–1.30 | normal | $\mu = 0$, $\sigma^2 = 10$ |
| PAV establishment | N addition (N) | 1.13 | 0.48–2.69 | normal | $\mu = 0$, $\sigma^2 = 10$ |
| PAV establishment | P addition (P) | 1.18 | 0.51–2.79 | normal | $\mu = 0$, $\sigma^2 = 10$ |
| PAV establishment | co-inoculation:N | 1.61 | 0.45–5.81 | normal | $\mu = 0$, $\sigma^2 = 10$ |
| PAV establishment | co-inoculation:P | 1.16 | 0.32–4.09 | normal | $\mu = 0$, $\sigma^2 = 10$ |
| PAV establishment | N:P | 0.65 | 0.20–2.17 | normal | $\mu = 0$, $\sigma^2 = 10$ |
| PAV establishment | co-inoculation:N:P | 0.81 | 0.14–4.78 | normal | $\mu = 0$, $\sigma^2 = 10$ |
| PAV establishment | random effect: time | 0.83 | 0.41–1.53 | Cauchy | $\chi_0 = 0$ $\gamma = 1$ |
| RPV establishment | intercept | 8.99 | 3.48–28.65 | normal | $\mu = 0$, $\sigma^2 = 10$ |
| RPV establishment | co-inoculation | 0.90 | 0.22–3.68 | normal | $\mu = 0$, $\sigma^2 = 10$ |
| RPV establishment | N addition (N) | 0.77 | 0.20–2.84 | normal | $\mu = 0$, $\sigma^2 = 10$ |
| RPV establishment | P addition (P) | 3.02 | 0.57–22.37 | normal | $\mu = 0$, $\sigma^2 = 10$ |
| RPV establishment | co-inoculation:N | 2.18 | 0.29–18.67 | normal | $\mu = 0$, $\sigma^2 = 10$ |
| RPV establishment | co-inoculation:P | 0.40 | 0.04–3.91 | normal | $\mu = 0$, $\sigma^2 = 10$ |
| RPV establishment | N:P | 0.22 | 0.02–1.72 | normal | $\mu = 0$, $\sigma^2 = 10$ |
| RPV establishment | co-inoculation:N:P | 1.96 | 0.10–44.42 | normal | $\mu = 0$, $\sigma^2 = 10$ |
| RPV establishment | random effect: time | 0.49 | 0.04–1.18 | Cauchy | $\chi_0 = 0$ $\gamma = 1$ |
| PAV ln(dens.) | intercept | 7.28 | 6.90–7.69 | normal | $\mu = 0$, $\sigma^2 = 10$ |
| PAV ln(dens.) | co-inoculation | 0.64* | 0.06–1.22 | normal | $\mu = 0.25$, $\sigma^2 = 0.73^\dagger$ |
| PAV ln(dens.) | N addition (N) | 0.17 | -0.36–0.69 | normal | $\mu = 0.08$, $\sigma^2 = 0.76^\dagger$ |
| PAV ln(dens.) | P addition (P) | -0.03 | -0.54–0.47 | normal | $\mu = -0.19$, $\sigma^2 = 0.83$ |
| PAV ln(dens.) | co-inoculation:N | -0.61 | -1.36–0.14 | normal | $\mu = 0$, $\sigma^2 = 1$ |
| PAV ln(dens.) | co-inoculation:P | -0.39 | -1.12–0.35 | normal | $\mu = 0$, $\sigma^2 = 1$ |
| PAV ln(dens.) | N:P | -0.01 | -0.72–0.70 | normal | $\mu = 0.17$, $\sigma^2 = 1.06$ |
| PAV ln(dens.) | co-inoculation:N:P | -0.04 | -1.01–0.93 | normal | $\mu = 0$, $\sigma^2 = 1$ |
| PAV ln(dens.) | sigma | 1.00 | 0.89–1.11 | Student's t | $\nu = 3$, $\hat{\mu} = 0$, $\hat{\sigma} = 10$ |
| PAV ln(dens.) | autocorrelation: time | 0.10 | -0.06–0.27 | bounded | lower = -1, upper = 1 |
| RPV ln(dens.) | intercept | 9.53 | 9.00–10.03 | normal | $\mu = 0$, $\sigma^2 = 10$ |
| RPV ln(dens.) | co-inoculation | 0.31 | -0.36–0.97 | normal | $\mu = -0.35$, $\sigma^2 = 0.66^\dagger$ |
| RPV ln(dens.) | N addition (N) | 0.03 | -0.60–0.67 | normal | $\mu = -0.21$, $\sigma^2 = 0.58^\dagger$ |
| RPV ln(dens.) | P addition (P) | 0.40 | -0.29–1.10 | normal | $\mu = 0.79$, $\sigma^2 = 0.97^\dagger$ |
| RPV ln(dens.) | co-inoculation:N | 0.33 | -0.58–1.21 | normal | $\mu = -0.54$, $\sigma^2 = 0.84^\dagger$ |
| RPV ln(dens.) | co-inoculation:P | -0.83 | -1.82–0.14 | normal | $\mu = -1.22$, $\sigma^2 = 1.39^\dagger$ |
| RPV ln(dens.) | N:P | 0.05 | -0.91–0.98 | normal | $\mu = -0.26$, $\sigma^2 = 1.15^\dagger$ |
| RPV ln(dens.) | co-inoculation:N:P | 0.51 | -0.84–1.85 | normal | $\mu = 0.74$, $\sigma^2 = 1.61$ |
| RPV ln(dens.) | sigma | 1.82 | 1.68–1.98 | Student's t | $\nu = 3$, $\hat{\mu} = 0$, $\hat{\sigma} = 10$ |
| RPV ln(dens.) | autocorrelation: time | 0.15 | 0.03–0.27 | bounded | lower = -1, upper = 1 |
| PAV transmission | intercept | 4.67 | 1.39–17.24 | normal | $\mu = 0$, $\sigma^2 = 10$ |
| PAV transmission | density | 0.82 | 0.52–1.28 | normal | $\mu = -0.22$, $\sigma^2 = 0.25^\dagger$ |
| PAV transmission | co-infection | 0.68 | 0.14–2.99 | normal | $\mu = 6.05$, $\sigma^2 = 4.13^\dagger$ |
| PAV transmission | N addition (source) | 1.72 | 0.72–4.15 | normal | $\mu = 0.64$, $\sigma^2 = 0.78^\dagger$ |
| PAV transmission | P addition (source) | 0.83 | 0.35–1.91 | normal | $\mu = -1.18$, $\sigma^2 = 0.71^\dagger$ |

| | | | | | |
|---|---|---|---|---|---|
| PAV transmission | N addition (recipient) | 0.75 | 0.20–2.83 | normal | $\mu = 0$, $\sigma^2 = 10$ |
| PAV transmission | P addition (recipient) | 0.18* | 0.05–0.57 | normal | $\mu = 0$, $\sigma^2 = 10$ |
| PAV transmission | $N_{source}$:$P_{source}$ | 0.79 | 0.24–2.53 | normal | $\mu = 2.07$, $\sigma^2 = 0.93^\dagger$ |
| PAV transmission | $N_{recipient}$:$P_{recipient}$ | 8.88* | 1.35–64.55 | normal | $\mu = 0$, $\sigma^2 = 10$ |
| PAV transmission | density:$N_{source}$ | 2.01 | 0.95–5.13 | normal | $\mu = 0$, $\sigma^2 = 10$ |
| PAV transmission | density:$P_{source}$ | 2.72 | 0.84–10.55 | normal | $\mu = 0$, $\sigma^2 = 10$ |
| PAV transmission | density:$N_{recipient}$ | 1.36 | 0.50–4.95 | normal | $\mu = 0$, $\sigma^2 = 10$ |
| PAV transmission | density:$P_{recipient}$ | 0.44* | 0.17–0.95 | normal | $\mu = 0$, $\sigma^2 = 10$ |
| PAV transmission | co-infection:$N_{source}$ | 0.22* | 0.05–0.93 | normal | $\mu = -7.67$, $\sigma^2 = 4.16^\dagger$ |
| PAV transmission | co-infection:$P_{source}$ | 0.63 | 0.14–2.72 | normal | $\mu = 0$, $\sigma^2 = 10$ |
| PAV transmission | co-infection:$N_{recipient}$ | 1.74 | 0.35–8.54 | normal | $\mu = 0$, $\sigma^2 = 10$ |
| PAV transmission | co-infection:$P_{recipient}$ | 8.60* | 2.00–41.27 | normal | $\mu = 0$, $\sigma^2 = 10$ |
| PAV transmission | density:$N_{source}$:$P_{source}$ | 0.13 | 0.01–1.50 | normal | $\mu = 0$, $\sigma^2 = 10$ |
| PAV transmission | density:$N_{recipient}$:$P_{recipient}$ | 7.24 | 0.88–110.44 | normal | $\mu = 0$, $\sigma^2 = 10$ |
| PAV transmission | co:$N_{source}$:$P_{source}$ | 1.71 | 0.27–11.02 | normal | $\mu = 0$, $\sigma^2 = 10$ |
| PAV transmission | co:$N_{recipient}$:$P_{recipient}$ | 0.17 | 0.02–1.58 | normal | $\mu = 0$, $\sigma^2 = 10$ |
| PAV transmission | random effect: time | 0.56 | 0.04–1.38 | Cauchy | $\chi_0 = 0$ $\gamma = 1$ |
| PAV transmission | random effect: block | 0.33 | 0.01–1.13 | Cauchy | $\chi_0 = 0$ $\gamma = 1$ |
| RPV transmission | intercept | 2.46 | 1.06–6.08 | normal | $\mu = 0$, $\sigma^2 = 10$ |
| RPV transmission | density | 1.30* | 1.03–1.63 | normal | $\mu = 0.20$, $\sigma^2 = 0.13^\dagger$ |
| RPV transmission | co-infection | 0.84 | 0.46–1.56 | normal | $\mu = -0.17$, $\sigma^2 = 0.45^\dagger$ |
| RPV transmission | N addition (source) | 0.92 | 0.54–1.60 | normal | $\mu = 0.64$, $\sigma^2 = 0.42^\dagger$ |
| RPV transmission | P addition (source) | 1.60 | 0.85–3.04 | normal | $\mu = 0.12$, $\sigma^2 = 0.70^\dagger$ |
| RPV transmission | N addition (recipient) | 2.81* | 1.25–6.69 | normal | $\mu = 0$, $\sigma^2 = 10$ |
| RPV transmission | P addition (recipient) | 1.33 | 0.64–2.86 | normal | $\mu = 0$, $\sigma^2 = 10$ |
| RPV transmission | $N_{source}$:$P_{source}$ | 0.66 | 0.29–1.49 | normal | $\mu = 0$, $\sigma^2 = 0.86^\dagger$ |
| RPV transmission | $N_{recipient}$:$P_{recipient}$ | 0.90 | 0.26–3.12 | normal | $\mu = 0$, $\sigma^2 = 10$ |
| RPV transmission | density:$N_{source}$ | 0.77 | 0.22–2.56 | normal | $\mu = 0$, $\sigma^2 = 10$ |
| RPV transmission | density:$P_{source}$ | 1.19 | 0.70–2.26 | normal | $\mu = 0$, $\sigma^2 = 10$ |
| RPV transmission | density:$N_{recipient}$ | 0.78 | 0.47–1.38 | normal | $\mu = 0$, $\sigma^2 = 10$ |
| RPV transmission | density:$P_{recipient}$ | 0.68 | 0.41–1.10 | normal | $\mu = 0$, $\sigma^2 = 10$ |
| RPV transmission | co-infection:$N_{source}$ | 1.87 | 0.90–3.85 | normal | $\mu = 0.31$, $\sigma^2 = 0.60^\dagger$ |
| RPV transmission | co-infection:$P_{source}$ | 0.82 | 0.37–1.79 | normal | $\mu = -0.69$, $\sigma^2 = 0.97^\dagger$ |
| RPV transmission | co-infection:$N_{recipient}$ | 0.87 | 0.29–2.60 | normal | $\mu = 0$, $\sigma^2 = 10$ |
| RPV transmission | co-infection:$P_{recipient}$ | 0.77 | 0.29–1.99 | normal | $\mu = 0$, $\sigma^2 = 10$ |
| RPV transmission | density:$N_{source}$:$P_{source}$ | 0.63 | 0.15–2.49 | normal | $\mu = 0$, $\sigma^2 = 10$ |
| RPV transmission | density:$N_{recipient}$:$P_{recipient}$ | 4.27* | 1.57–15.61 | normal | $\mu = 0$, $\sigma^2 = 10$ |
| RPV transmission | co:$N_{source}$:$P_{source}$ | 1.21 | 0.41–3.66 | normal | $\mu = -0.01$, $\sigma^2 = 1.14^\dagger$ |
| RPV transmission | co:$N_{recipient}$:$P_{recipient}$ | 0.81 | 0.16–4.16 | normal | $\mu = 0$, $\sigma^2 = 10$ |
| RPV transmission | random effect: time | 0.30 | 0.02–0.75 | Cauchy | $\chi_0 = 0$ $\gamma = 1$ |
| RPV transmission | random effect: block | 0.53 | 0.11–1.47 | Cauchy | $\chi_0 = 0$ $\gamma = 1$ |

Notes: All predictors had an r-hat of 1.00. Estimates for establishment and transmission are odds ratios. Estimates for random effects are standard deviation of the intercept. Density was centered and scaled when it was a predictor. Asterisk indicates estimate has 95% CI (omitting intercepts and random effects) that do not include zero (for log-transformed density) or one (for establishment and transmission), which suggests that "no effect" is absent from the most probable estimate values. $^\dagger$Indicates informative priors based on data associated with Lacroix et al. (2017). CI = credible interval, prior dist. = prior distribution, hyperparameters = parameters of prior distribution.

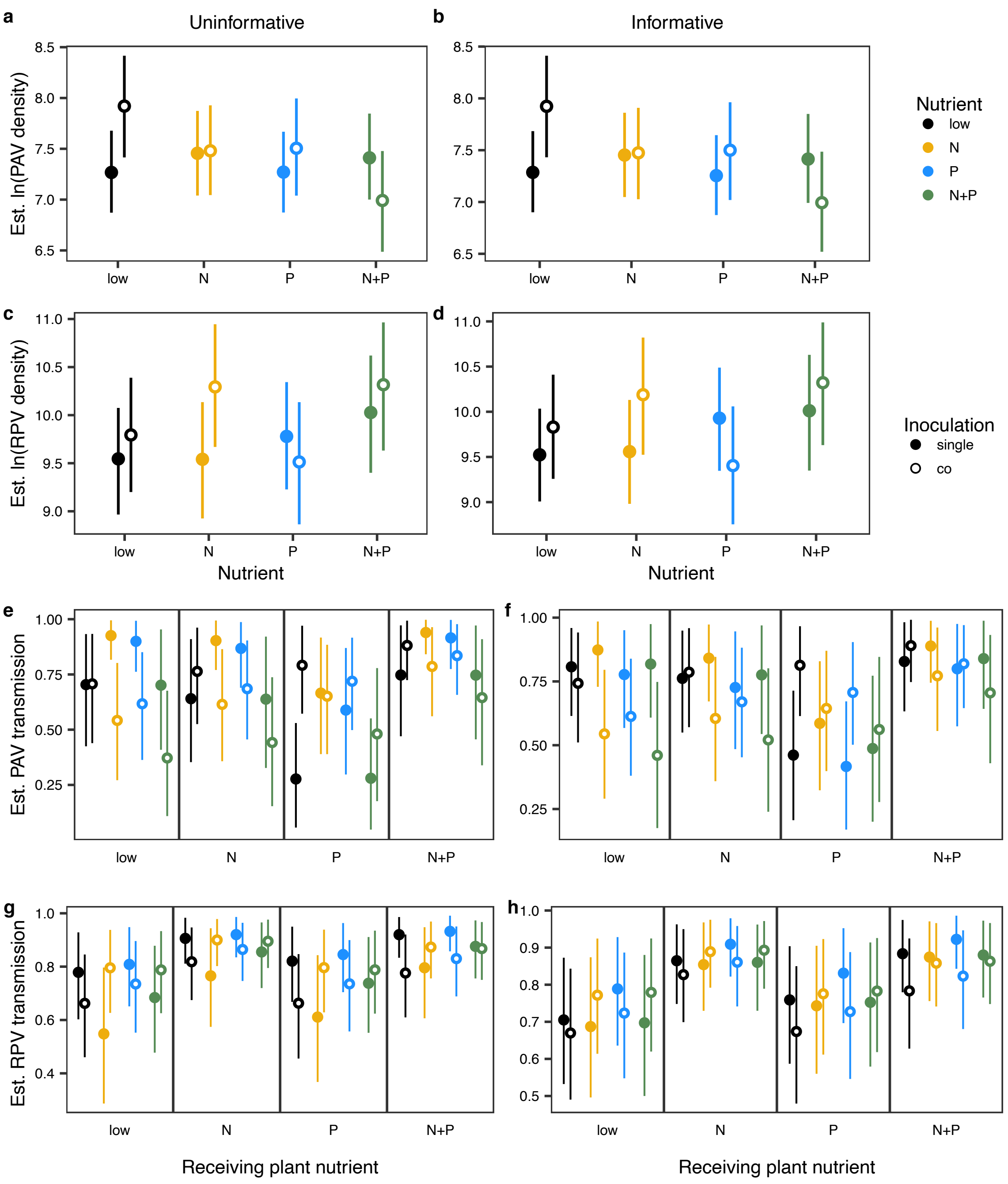


**Appendix S2: Figure S1**. The model estimated mean and 95% credible intervals of log-transformed (a–b) PAV and (c–d) RPV density and the transmission of (e–f) PAV and (g–h) RPV for each inoculation and nutrient treatment, pooled over all time points. Models have (a, c, e, g) uninformative priors or (b, d, f, h) informative priors (Appendix S2: Table S1), but are otherwise equivalent. Panels with informative priors (b, d, f, h) are the same as the corresponding panels in Fig. 1 and 2.

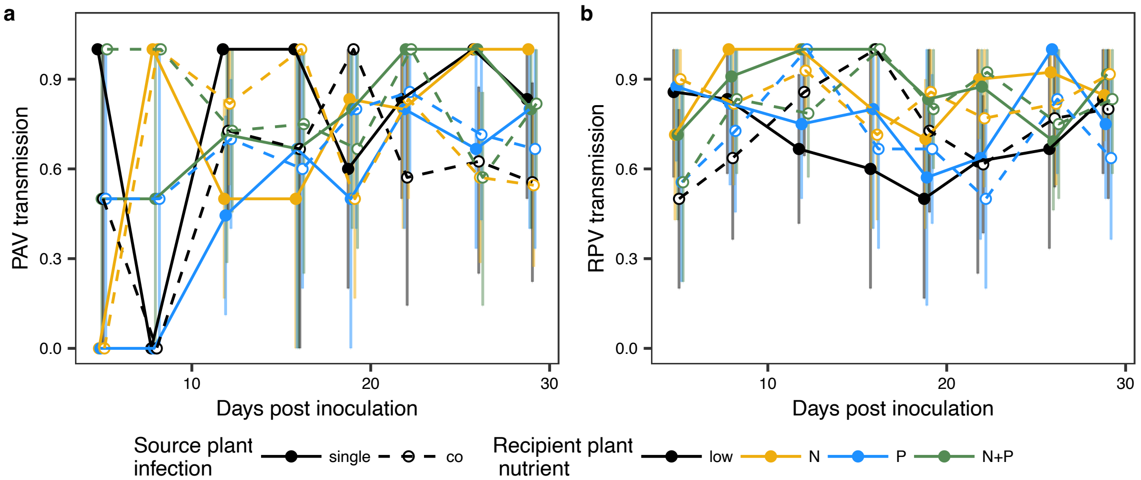


**Appendix S2: Figure S2.** The effects of nutrient addition to the recipient plant (N, P, or both) and co-infection of the source plant on (a) PAV and (b) RPV transmission (proportion of recipient plants infected) measured over time (mean ± 95% nonparametric bootstrap confidence intervals, values are averaged across source plant nutrient treatments).

**Appendix S2 literature cited**

Lacroix, C., E. W. Seabloom, and E. T. Borer. 2017. Environmental nutrient supply directly alters plant traits but indirectly determines virus growth rate. Frontiers in Microbiology 8:2116.
