## Appendix S3 for "Host nutrition mediates interactions between plant viruses, altering transmission and predicted disease spread"

**Appendix S3: Extended methods and results for mathematical model**
"Host nutrition mediates interactions between plant viruses, altering transmission and predicted disease spread"
Amy E. Kendig, Elizabeth T. Borer, Emily N. Boak, Tashina C. Picard, and Eric W. Seabloom
*Ecology*

The mathematical model includes several assumptions. First, transmission is assumed to be frequency-dependent (i.e., terms are divided by the total population size, $N$), which is a common assumption for vector-borne disease transmission (Keeling and Rohani 2008). Contact between susceptible and infected hosts in the model encompasses virus acquisition by the aphid and inoculation of the susceptible plant. We chose to omit state variables representing vectors from the model (e.g., Mordecai et al. 2016) because our experiment does not provide the information required to estimate separate values for vector acquisition and inoculation. Because the experiment included forced contact between susceptible hosts and vectors, we did not estimate the effects of host nutrition or infection status on contact rates. We omitted contact rates from the transmission values, which likely leads to overestimates of transmission relative to actual values. However, we are primarily interested in the relative effects of nutrient addition on infection prevalence, not the absolute values of infection prevalence. We assumed that co-infected plants can transmit both pathogens ($q_P\beta_P q_R\beta_R I_C$) or one of the two pathogens (e.g., $q_P\beta_P(1 - q_R\beta_R)I_C$). The value $1 - q_i\beta_i$, where $i$ is $P$ or $R$, is constrained to be between (and including) zero and one due to the methods we used to estimate $q_i$ and $\beta_i$. Plants can also become sequentially infected (e.g., $\beta_P(I_P + q_P I_C)I_R/N$), but the probability of a virus establishing in a plant that is already infected is equal to establishment in a completely susceptible plant. We assumed that no new plants germinate, that no plants die from infection or otherwise, and that all infections persist through the growing season, which is the length of the simulation.

To assign values to $\beta_P$, $\beta_R$, $q_P$, and $q_R$, we first extracted 15000 posterior samples of each predictor variable coefficient estimated with the empirical transmission model (Eq. 1). We assumed that transmission only occurred between source and recipient plants grown with the same nutrient treatment and estimated separate parameter sets for each infection-nutrient combination for each virus (Appendix S3: Table S1). We set the predictor variables in Eq. 1 to reflect each infection-nutrient combination, substituting one or zero for the binary variables and the average virus density for the infection-nutrient combination for "density". We then calculated transmission using each of 15000 posterior samples and took the average across all samples. We used the average transmission from singly infected plants for values of $\beta_P$ and $\beta_R$ and divided transmission from co-infected plants by transmission from singly infected plants to determine values for $q_P$ and $q_R$ ("all processes", Appendix S3: Table S1). We repeated these calculates for two more scenarios (Appendix S3: Table S2), in which only predictor variables related to density were included ("virus density–dependent") and in which only predictor variables unrelated to density were included ("virus density–independent"). Because we allowed aphids four days for acquisition and two days for inoculation in the experiment, we assumed that the parameter values represent a six-day transmission process and conducted simulations for 20 time-steps to represent 120 days. Simulations were repeated for each nutrient treatment within each scenario.

**Appendix S3: Table S1.** Mathematical model parameters

| Parameter | Meaning | Nutrient | All processes | | Virus density–dependent | | Virus density–independent | |
|---|---|---|---|---|---|---|---|---|
| | | | Mean | 95% HDI | Mean | 95% HDI | Mean | 95% HDI |
| $\beta_P$ | transmission of PAV from singly infected plants | Low | 0.82 | (0.63–0.96) | 0.82 | (0.63–0.96) | 0.81 | (0.61–0.96) |
| | | N | 0.80 | (0.6–0.95) | 0.75 | (0.53–0.95) | 0.84 | (0.67–0.97) |
| | | P | 0.42 | (0.17–0.66) | 0.81 | (0.62–0.96) | 0.42 | (0.17–0.67) |
| | | N + P | 0.80 | (0.6–0.97) | 0.74 | (0.48–0.97) | 0.84 | (0.64–0.99) |
| $\beta_R$ | transmission of RPV from singly infected plants | Low | 0.70 | (0.52–0.87) | 0.70 | (0.52–0.87) | 0.71 | (0.53–0.87) |
| | | N | 0.86 | (0.75–0.97) | 0.72 | (0.53–0.88) | 0.85 | (0.73–0.97) |
| | | P | 0.83 | (0.7–0.95) | 0.70 | (0.52–0.87) | 0.83 | (0.7–0.95) |
| | | N + P | 0.87 | (0.75–0.97) | 0.69 | (0.51–0.86) | 0.88 | (0.77–0.97) |
| $q_P$ | modification of transmission for PAV from co-infected plants | Low | 0.90 | (0.57–1.22) | 0.97 | (0.89–1.04) | 0.93 | (0.59–1.26) |
| | | N | 0.91 | (0.54–1.29) | 1.18 | (0.95–1.52) | 0.72 | (0.42–1.01) |
| | | P | 1.86 | (0.86–3.16) | 1.00 | (0.96–1.04) | 1.88 | (0.86–3.24) |
| | | N + P | 0.82 | (0.47–1.15) | 1.01 | (0.99–1.05) | 0.84 | (0.51–1.13) |
| $q_R$ | modification of transmission for RPV from co-infected plants | Low | 0.99 | (0.79–1.19) | 1.03 | (1–1.07) | 0.95 | (0.77–1.16) |
| | | N | 1.03 | (0.9–1.19) | 0.98 | (0.86–1.1) | 1.05 | (0.89–1.22) |
| | | P | 0.88 | (0.68–1.07) | 1.01 | (0.94–1.08) | 0.88 | (0.67–1.06) |
| | | N + P | 1.01 | (0.86–1.18) | 1.08 | (0.94–1.25) | 0.98 | (0.83–1.13) |

Notes: For more information about the scenarios, see Appendix S3: Table S2.

**Appendix S3: Table S2.** Scenarios for estimating mathematical model parameters

| Scenario | Virus density value | Equation 1 parameters |
|---|---|---|
| Virus density–dependent and independent transmission (all processes) | Average for infection-nutrient combination | intercept + density + $N_{source}$:density + $P_{source}$:density + $N_{recipient}$:density + $P_{recipient}$:density + $N_{source}$: $P_{source}$:density + $N_{recipient}$: $P_{recipient}$:density + co-infection × ($N_{source}$ × $P_{source}$ + $N_{recipient}$ × $P_{recipient}$) (i.e., all of the parameters) |
| Virus density–dependent transmission | Average for infection-nutrient combination | intercept + density + $N_{source}$:density + $P_{source}$:density + $N_{recipient}$:density + $P_{recipient}$:density + $N_{source}$: $P_{source}$:density + $N_{recipient}$: $P_{recipient}$:density |
| Virus density–independent transmission | Overall average | intercept + co-infection × ($N_{source}$ × $P_{source}$ + $N_{recipient}$ × $P_{recipient}$) |

Notes: Equation 1 refers to the main text and the transmission models in Appendix S2: Table S1. The symbol "×" indicates that variables on either side are included as main effects and interactions.

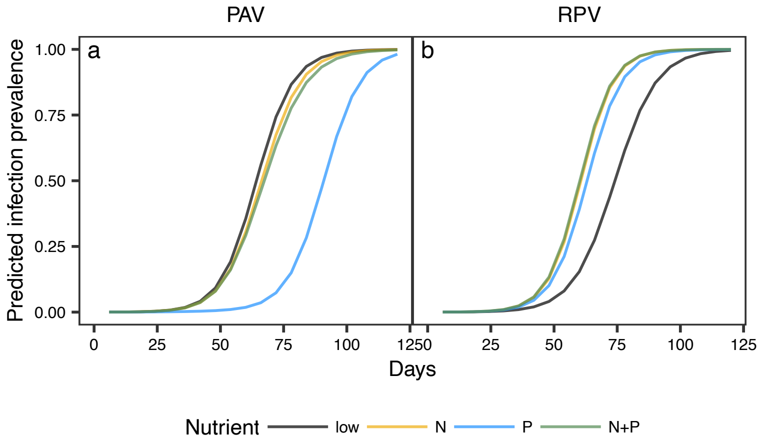


**Appendix S3: Figure S1.** The predicted infection prevalence of (a) PAV and (b) RPV when virus density–dependent and independent processes affect transmission (i.e., “all processes” in Fig. 4 and Appendix S3: Table S2) and prevalence is summed over singly and co-infected hosts. Values are estimated from Eq. 2. The line for “N” is under “N+P” in panel b.

### Appendix S3 literature cited

Keeling, M. J., and P. Rohani. 2008. Modeling Infectious Diseases in Humans and Animals. Princeton University Press, Princeton, NJ.

Mordecai, E. A., K. Gross, and C. E. Mitchell. 2016. Within-host niche differences and fitness trade-offs promote coexistence of plant viruses. The American Naturalist 187:E13–E26.
